# Molecular evolution of the mammalian kinetochore complex

**DOI:** 10.1101/2024.06.27.600994

**Authors:** Uma P. Arora, Beth L. Dumont

## Abstract

Mammalian centromeres are satellite-rich chromatin domains that serve as sites for kinetochore complex assembly. Centromeres are highly variable in sequence and satellite organization across species, but the processes that govern the co-evolutionary dynamics between rapidly evolving centromeres and their associated kinetochore proteins remain poorly understood. Here, we pursue a course of phylogenetic analyses to investigate the molecular evolution of the complete kinetochore complex across primate and rodent species with divergent centromere repeat sequences and features. We show that many protein components of the core centromere associated network (CCAN) harbor signals of adaptive evolution, consistent with their intimate association with centromere satellite DNA and roles in the stability and recruitment of additional kinetochore proteins. Surprisingly, CCAN and outer kinetochore proteins exhibit comparable rates of adaptive divergence, suggesting that changes in centromere DNA can ripple across the kinetochore to drive adaptive protein evolution within distant domains of the complex. Our work further identifies kinetochore proteins subject to lineage-specific adaptive evolution, including rapidly evolving proteins in species with centromere satellites characterized by higher-order repeat structure and lacking CENP-B boxes. Thus, features of centromeric chromatin beyond the linear DNA sequence may drive selection on kinetochore proteins. Overall, our work spotlights adaptively evolving proteins with diverse centromere-associated functions, including centromere chromatin structure, kinetochore protein assembly, kinetochore-microtubule association, cohesion maintenance, and DNA damage response pathways. These adaptively evolving kinetochore protein candidates present compelling opportunities for future functional investigations exploring how their concerted changes with centromere DNA ensure the maintenance of genome stability.

## Introduction

The kinetochore is a large, multiprotein complex that orchestrates chromosome segregation to assure the faithful distribution of a replicated genome from a mother cell to its daughter cells. The kinetochore complex also functions as a mechanosensory structure that corrects errors in microtubule attachment and regulates the transition from metaphase to anaphase. Disruptions to kinetochore complex function can lead to chromosome mis-segregation or cell cycle arrest, outcomes with adverse impacts on organismal development, survival, and function (McKinley and Cheeseman 2016).

Despite their crucial biological functions, centromeres are among the most rapidly evolving loci in mammalian genomes. In many species, centromeres are comprised of repetitive satellite DNA subject to exceptionally high rates of mutation through replication slippage and unequal crossing over (Charlesworth, et al. 1994; Ugarković and Plohl 2002; Logsdon, et al. 2024). Rapid centromere DNA evolution is likely further fueled by centromere drive, a phenomenon characterized by the biased segregation of chromosomes with a “stronger” centromere into the functional oocyte during asymmetric female meiosis (Chmátal, et al. 2014; Iwata-Otsubo, et al. 2017; Akera, et al. 2019). In this manner, centromere drive mimics the action of intense positive selection and leads to the rapid fixation of stronger centromere variants. Together, mutation and drive are hypothesized to underlie the remarkable size and sequence variability in centromere satellites between species (Alkan, et al. 2011; Melters, et al. 2013; Logsdon, et al. 2024).

The selfish transmission of alleles via centromere drive is theorized to pose fitness costs (Henikoff, et al. 2001; Malik 2009). Although the nature of this fitness cost is not well understood, centromere drive imposes selection for the emergence of suppressive mechanisms that restore segregation balance. Adaptive changes to protein components of the kinetochore complex may counter the action of selfish centromere variants to restore the parity of chromosome segregation (Henikoff, et al. 2001; Malik 2009). Consistent with this hypothesis, multiple kinetochore proteins are adaptively co-evolving with centromere DNA across diverse eukaryotic taxa (Talbert, et al. 2004; Schueler, et al. 2010; Drinnenberg, et al. 2016; van Hooff, et al. 2017; Tromer, et al. 2019; Kumon, et al. 2021). In particular, prior analyses have uncovered signals of adaptive evolution in proteins of the core centromere associated network (CCAN), including CENP-A, CENP-B, and CENP-C (Malik and Henikoff 2001; Malik, et al. 2002; Talbert, et al. 2004; Schueler, et al. 2010). CCAN proteins intimately interact with centromere DNA throughout the cell cycle and are potentially most vulnerable to centromere DNA sequence changes.

Kinetochore proteins function in complex interactive networks, with individual kinetochore proteins capable of influencing the recruitment, assembly, and function of other kinetochore proteins (Beck and Llopart 2015). Thus, the co-evolutionary response of kinetochore proteins to sequence changes in centromere DNA could extend beyond CCAN components, rippling to other interacting and functionally related proteins in the kinetochore (Beck and Llopart 2015). This insight emphasizes the crucial need for holistic investigations into the molecular evolution of the full kinetochore complex, inclusive of all its protein components. However, few studies have employed such a comprehensive approach, and of those, most have adopted a very broad phylogenetic lens (Meraldi, et al. 2006; van Hooff, et al. 2017). Examination of kinetochore proteins across eukaryotes with vastly different centromere DNA architectures has led to the discovery that some kinetochore constituents are broadly conserved between higher eukaryotes and yeast, whereas other kinetochore components have undergone significant diversification. For example, *Drosophila melanogaster* and *Caenorhabditis elegans* lack most CCAN components (van Hooff, et al. 2017). Additionally, several spindle assembly checkpoint proteins (BUB1/BUBR1/MAD3) have been duplicated and sub-functionalized multiple times during eukaryotic evolution (van Hooff, et al. 2017). These broad evolutionary surveys have exposed the dynamic molecular composition of the kinetochore, but the mechanisms of kinetochore evolution across organisms with more similar centromere DNA architectures and identical casts of kinetochore proteins are less well understood.

To address this knowledge gap, recent studies have examined the molecular evolution of many kinetochore proteins on finer phylogenetic timescales, revealing new evolutionary insights. For example, Kumon et al. showed that, relative to other proteins, kinetochore proteins exhibit high rates of adaptive evolution in rodents (Kumon, et al. 2021). Intriguingly, despite the lack of direct contact with centromere satellite DNA, microtubule destabilizer recruitment proteins are evolving via positive selection in mammals (Kumon, et al. 2021; Pontremoli, et al. 2021). This observation lends support to the hypothesis that rapid centromere DNA sequence evolution exerts pervasive selection pressures on diverse kinetochore components and reinforces the need for investigations that interrogate mechanisms of evolution across all proteins in this complex.

The satellite DNA composition of mammalian centromeres varies widely with respect to repeat size, heterogeneity, and structure (Melters et al. 2013). Whether basic architectural features of centromere repeat organization influence the regime or intensity of adaptive kinetochore protein evolution remains an open question. Investigating adaptive evolution in a group of closely related mammals with diverse regional centromere features could uncover general rules of kinetochore protein evolution governed by specific aspects of centromere architecture.

One notable architectural feature that varies across rodent and primate centromeres is the presence and abundance of the CENP-B box. The CENP-B box is a canonical 17-bp motif embedded within centromere satellite repeat units that enables sequence-specific binding of CENP-B, a CCAN component. Through interactions with CENP-A and CENP-C, CENP-B stabilizes the kinetochore (Suzuki, et al. 2004; Fachinetti, et al. 2015; Fujita, et al. 2015), although CENP-B is not required for the maintenance of established centromeres. Notably, there are no CENP-B boxes on human and mouse Y-chromosome centromere satellite repeats (Masumoto, et al. 1989; Earnshaw, et al. 1991) and CENP-B knockout mice are viable (Hudson, et al. 1998; Perez-Castro, et al. 1998). Further, not all species have centromere satellites that contain CENP-B boxes, even though functional CENP-B protein is found at centromeres of species lacking the sequence motif (Kipling, et al. 1995; Goldberg, et al. 1996). The functional impact of the evolutionary changes in CENP-B box occurrence is unclear.

A second variable characteristic of primate and rodent centromeres is higher-order repeat (HOR) structure. HORs are tandem arrays of large repeat units individually composed of repetitions of smaller repeat units. HOR structure is commonly present at primate centromeres, but rarely defines rodent centromeres. Exploration is needed to assess whether this feature of satellite repeat organization shapes kinetochore protein organization and function via adaptive evolution (Wong and Rattner 1988; Melters, et al. 2013; Gambogi, et al. 2023).

Here, we profile a subset of mammals with a conserved cast of kinetochore proteins but that vary with respect to multiple regional centromere features. We employ phylogenetic approaches to model distinct evolutionary regimes defined by centromere attributes like HOR structure and CENP-B box occurrence. Our ensemble approach investigates all annotated kinetochore proteins to enable unbiased discovery of adaptive evolution in individual protein components, as well as co-evolutionary trends between proteins. Our work uncovers specific kinetochore proteins evolving under distinct adaptive regimes and nominates candidates for future functional studies aiming to directly assess the functional consequences of kinetochore protein and centromere DNA co-evolution.

## RESULTS

### Differential rates of kinetochore protein evolution across primates and rodents

We set out to investigate evidence for adaptive evolution of kinetochore complex proteins across 17 mammalian species that exhibit differences in various aspects of centromere satellite repeat sequence and organization (Figure 1A). We selected species within the Euarchontoglires superorder (rodents and primates) with high-quality and well-annotated whole genome sequences. For each species, protein coding sequences were obtained from 215 genes with GO term associations that contained the word “centromere” or “kinetochore” (Supplementary Table 1). These proteins execute diverse cellular and biological functions, including association with centromere DNA, kinetochore protein complex assembly, microtubule binding, and mechanosensory and checkpoint pathway signaling.

**Figure 1:**
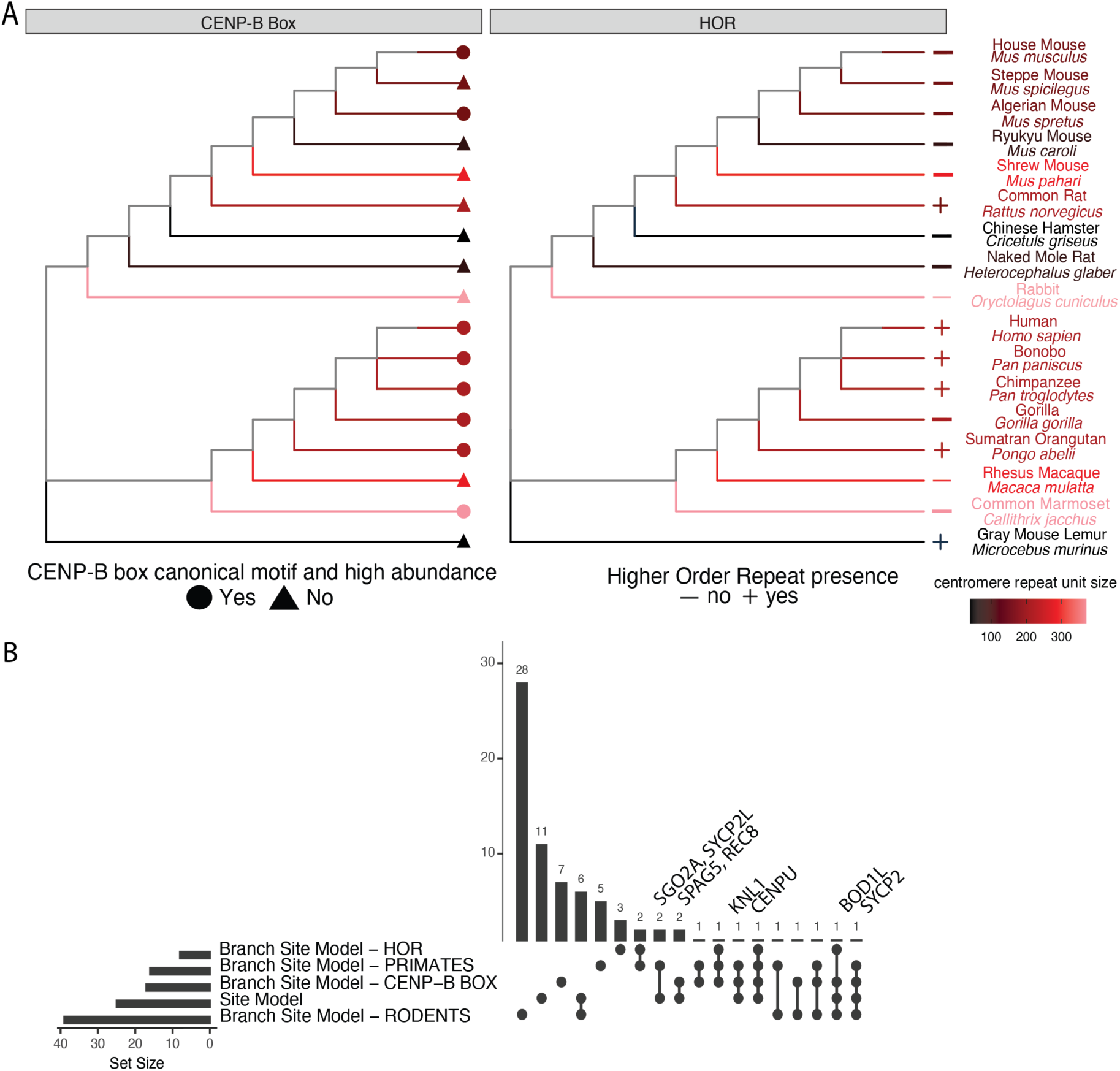
Phylogenetic analysis of adaptive evolution across the kinetochore complex. (A) Cladogram representing the phylogenetic relationships among the 17 rodent and primate species selected for analysis. Trees are annotated according to the presence/absence of CENP-B box motifs in consensus centromere satellite sequences (left) and higher-order repeat structure (right). (B) Upset plot displaying the number of adaptively evolving kinetochore proteins under the site and branch-site models. The horizontal bars represent the number of proteins tested in each of the models, and the vertical bars represent the number of proteins that are significant under the specified subset of models, as indicated by the dots below.

We used PAML (v 4.8; (Yang 1997, 2007)) to estimate the ratio of non-synonymous (dN) to synonymous (dS) amino acid substitutions, *ω*, at each kinetochore protein under several different models (see Methods), each invoking a distinct number of evolutionary rate classes. The estimate of *ω* under the best fit model (M2) for the subset of kinetochore proteins exhibiting significant signals of adaptive evolution is provided in Table 1.

**Table 1:**
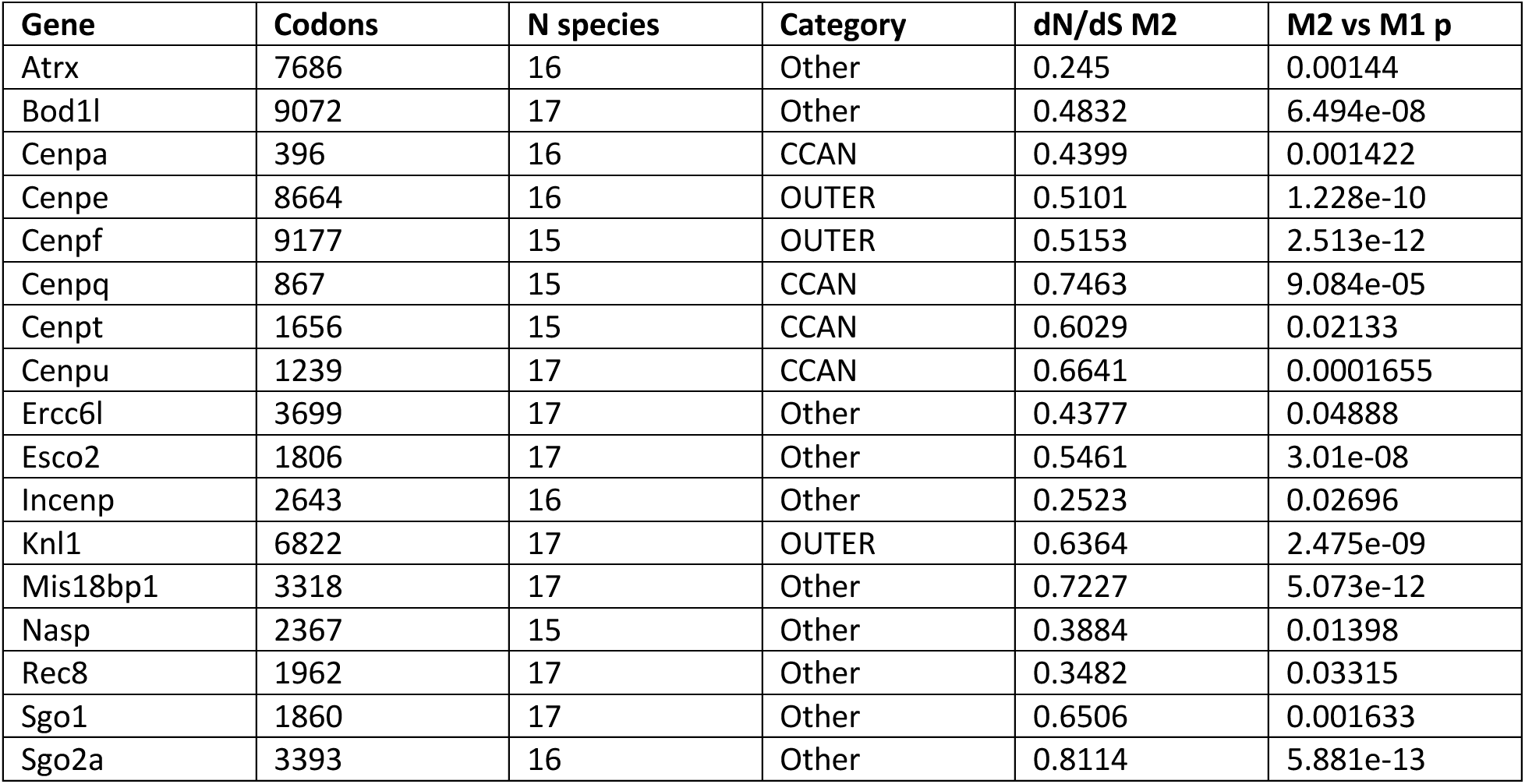
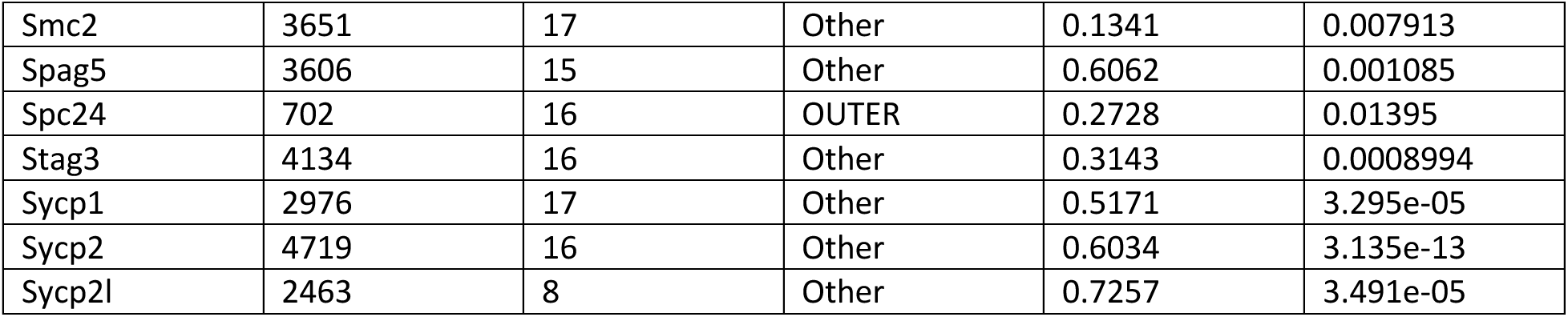
Kinetochore proteins with signals of adaptive evolution under the site model.

All kinetochore proteins have *ω* < 1 (Supplementary Figure 1A and B), consistent with the action of purifying selection across the primate and rodent clades. However, there is significant variation in *ω* across proteins, implying differences in the levels of constraint and/or the strength of adaptive evolution across kinetochore components (*ω*; range 0.00 - 0.81). Specifically, 13 proteins exhibit *ω* > 2 standard deviations (SD) greater than the mean for all kinetochore proteins (Brca2, Cenpc1, Cenpq, Cenpt, Cenpu, Hjurp, Itgb3bp, Knl1, Mis18bp1, Sgo1, Sgo2a, Sycp2, Sycp2l), consistent with rapid evolution via positive selection. At the other extreme, 29 proteins (Anapc16, Becn1, Bub3, Cbx1, Cbx3, Csnk1a1, Ctcf, Ctnnb1, Dapk3, Dctn5, Dynll1, H3c1, H3f3a, Ik, Kat8, Meaf6, Nudcd2, Pafah1b1, Peli1, Ppp2ca, Ppp2cb, Ppp2r1a, Racgap1, Septin7, Smc1a, Smc3, Stag2, Xpo1, Zfp207) have *ω* < 1 SD below the mean of all kinetochore proteins, suggesting a high level of functional constraint. Included among these are two histone H3 proteins (H3c1 and H3f3a) which have been previously shown to be under strong purifying selection (Malik and Henikoff 2003).

### Site-level evidence for adaptive kinetochore protein evolution

Theoretically, a protein evolving via positive selection is expected to have *ω* > 1. However, in practice, protein-wide estimates of *ω* rarely exceed this threshold as only a fraction of amino acid sites are subject to strong positive selection. Indeed, *ω* was less than one for all the kinetochore proteins surveyed in our analysis, motivating us to consider more sensitive approaches for detecting phylogenetic signals of adaptive protein evolution.

To this end, we performed likelihood-based statistical comparisons between nested site models to formerly test the hypothesis that a subset of sites in each kinetochore protein is evolving via positive selection across primates and rodents. We consider two sets of nested model comparisons. In the first comparison, we contrast a null, neutral model (M1) that features two classes of sites evolving under adaptive constraint (*ω* < 1) and neutrality (*ω* = 1) with a second model (M2) that includes a third class of sites evolving via positive selection (*ω* > 1). In the second comparison, we invoke more complex phylogenetic models that employ a larger number of site classes drawn from a beta distribution (M7 vs M8; n = 10 and 11 site classes, respectively). Overall, *ω* values are strongly positively correlated between the M2 and M8 models (Pearson’s correlation coefficient = 0.99, P < 2.2x10^-16^; Supplementary Figure 1C), assuring that our findings and conclusions are robust to model choice.

We reject the null model of neutral evolution for 25 and 71 kinetochore proteins under the M2 and M8 frameworks, respectively (11.6% and 33% of kinetochore proteins, respectively). As expected, proteins harboring signals of adaptive evolution (Figure 1B and Table 1) are marked by overall elevated *ω* compared to kinetochore proteins for which the null model cannot be rejected (Figure 2A; Wilcox rank sum test, P<1.1x10^-11^). All proteins identified as evolving under positive selection in the M2 framework are also detected by the M8 framework. Given their increased stringency, we focus on the M2 model results from here forward.

**Figure 2:**
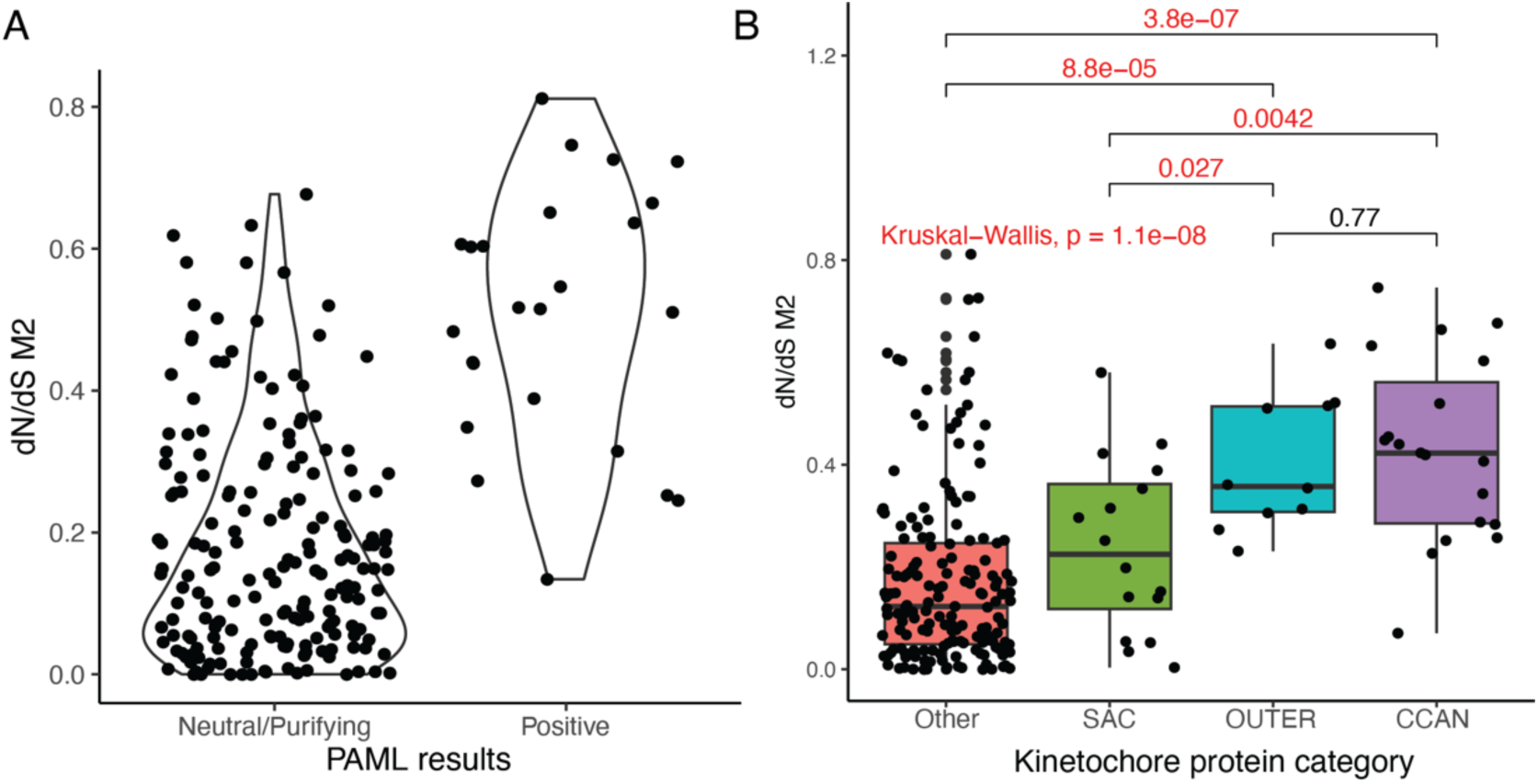
Site model analysis of adaptive kinetochore protein evolution. (A) Kinetochore proteins for which the null M1 site model is rejected harbor elevated dN/dS (*ω*) ratios compared to proteins for which the M1 model cannot be rejected (p < 2x10^-16^). (B) Boxplot representing the dN/dS (*ω*) ratios (y-axis) of kinetochore proteins based on their position and function in the kinetochore (x-axis). CCAN and outer kinetochore proteins have significantly higher dN/dS (*ω*) ratios compared to the SAC and other kinetochore proteins.

### Rate of protein evolution varies by functional role and physical position in the kinetochore complex

We next sought to determine whether the rate of protein evolution varies across different functional components of the kinetochore complex. We assigned kinetochore proteins to one of three categories: (1) proteins of the constitutive centromere associated network (CCAN) that closely associate with centromere DNA, (2) outer kinetochore proteins that associate with microtubules, and (3) spindle assembly checkpoint proteins (SAC). We predicted that CCAN proteins would experience, on average, higher rates of evolution than proteins in other functional categories due to their direct and sustained interactions with rapidly evolving centromere DNA.

Intriguingly, CCAN and outer kinetochore proteins exhibit comparable rates of evolution, and both groups of proteins have significantly higher values of *ω* compared to the SAC proteins (Figure 2B; Kruskal-Wallis, p<2.4x10^-9^, df = 5). Although not significant, a greater proportion of outer kinetochore proteins show evidence for adaptive evolution in primates and rodents (4/10; 40%), compared to the proportion of adaptively evolving CCAN proteins (4/19; 21%; FET p = 0.3904; Table 2).

### Weak correlation between species-level adaptive evolution and population-level diversity in kinetochore genes

The discovery of kinetochore proteins with signals of adaptive evolution across primates and rodents raises the question of whether kinetochore proteins also exhibit exceptional patterns of diversity at the population level. We quantified average pairwise sequence divergence (*π*) using single nucleotide polymorphisms (SNPs) along the gene sequence of each kinetochore protein in human and wild mouse populations (see Methods). We observed no difference in *π* between adaptively and non-adaptively evolving kinetochore genes identified under the site model in populations from either species (Figure 3A). Although the correlation between *π* and dN/dS (*ω*) in both species is statistically significant, the overall strength of the relationship between the level of polymorphism in a gene and the rate of adaptive divergence between species is weak (Figure 3B).

**Figure 3:**
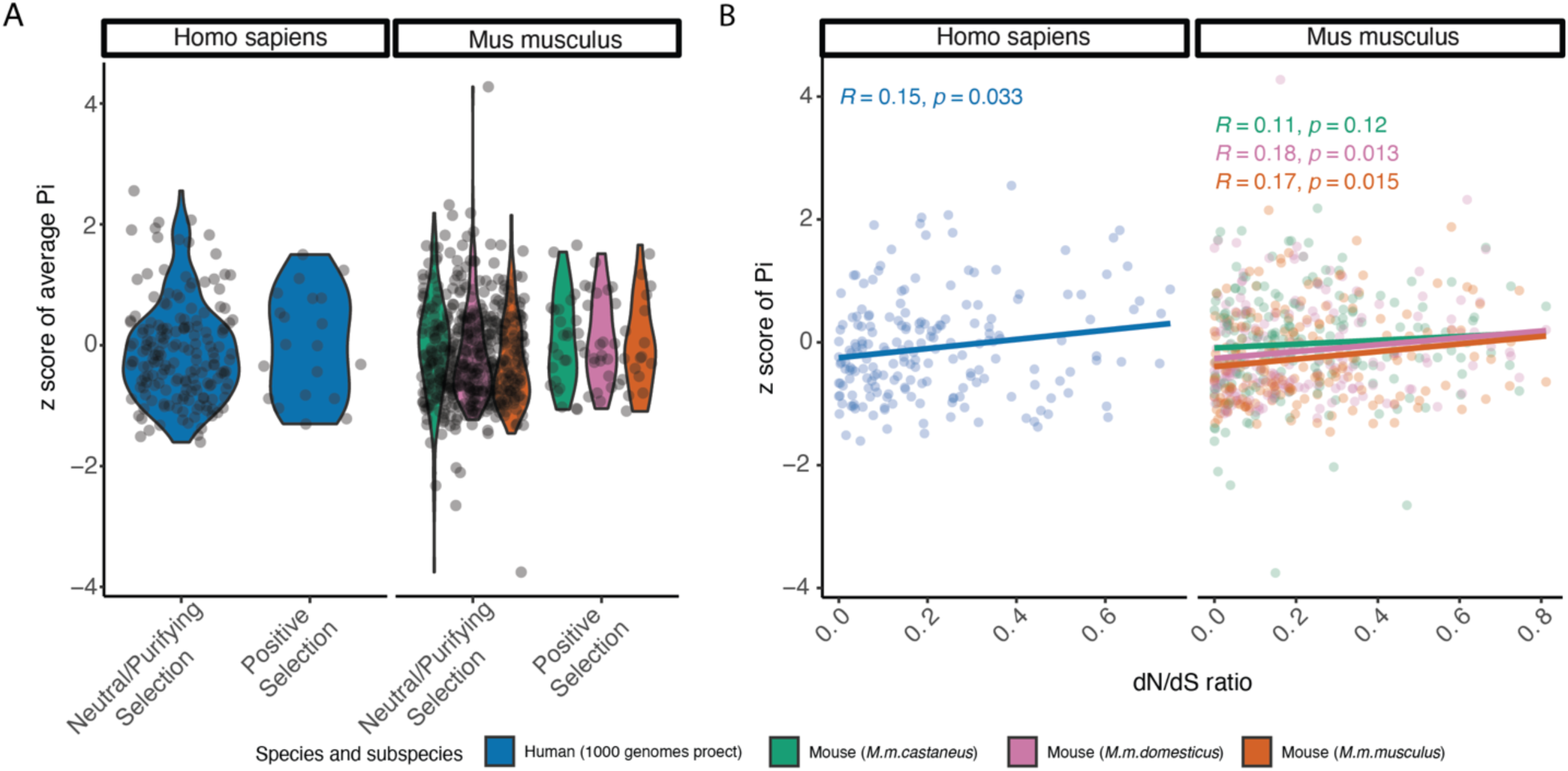
Relationship between adaptive protein divergence and population level diversity. (A) Distribution of normalized pairwise sequence divergence, *π*, and mode of evolution as assessed by the site model in humans and mice. (B) Correlation between dN/dS (*ω*) value of each gene under the site model and average pairwise sequence divergence (*π*). For both (A) and (B), human *π* values were computed from 1000 Genomes Project data, aggregating across all populations. Mouse data are from Harr et al. 2016 and partitioned by subspecies.

Next, we asked whether proteins that are diverging between species have been highlighted in published population genome-wide scans for recent positive selection. Proteins ascribed to the GO term “Kinetochore” are significantly enriched among genes within regions under recent positive selection in a wild-caught population of house mice from Taiwan (*M. m. castaneus*) (Lawal, et al. 2021), including *Mad1l1* (SAC)*, Ckap5, Ppp1r12a, Sec13, and Pbrm1*. In humans, *POGZ, CENP-O* (CCAN)*, DCTN1, SIN3A, HIRA, PPP1R12A,* and *DYNLT3* reside in genomic regions under recent positive selection (Sabeti, et al. 2007; Grossman, et al. 2013). However, none of the kinetochore genes under recent positive selection in mice or human populations are also identified as targets of long-term positive selection across primates and rodents. Thus, distinct selective forces appear to shape patterns of population variation and species divergence at kinetochore proteins.

### Branch-site model identifies sites co-evolving with specific centromere DNA features

Our analyses have spotlighted multiple kinetochore proteins that are adaptively evolving across Euarchontoglires, but may be underpowered to detect signals of positive selection operating on only a subset of lineages. Further, identifying cases of positive selection specific to species with shared centromere-related features could shed light on possible mechanistic drivers of adaptive evolution. To this end, we used the branch-site evolutionary modeling functionality in PAML to identify amino acid sites that are adaptively evolving across one or more lineages that share specific centromere features (Yang 1997, 2007). Specifically, we defined four different phylogenetic groupings for branch-site model analysis: rodents, primates, species harboring centromeres with a high abundance of cannonical CENP-B boxes, and species with HOR centromere satellite structure (Figure 1A). A total of 64 proteins are adaptively evolving under the branch-site model across one or more of these tested phylogenetic regimes (Figure 1B). Eight proteins exhibit site-specific *ω* rates consistent with adaptive evolution in species with HOR centromere architectures. We identify 16 and 39 proteins that show signals consistent with adaptive evolution along the primate and rodent lineages, respectively. A total of 17 proteins harbor signals of adaptive evolution specific to taxa with a high abundance of CENP-B boxes at centromeres. Of the proteins identified in this branch-site analysis, 21 (32%) have sites that are adaptively evolving under 2 or more of the tested regimes.

As in the site model analysis, we assessed whether genes harboring signals of clade-specific adaptive evolution also exhibit distinct patterns of population-level diversity. We again use nucleotide diversity (*π*) estimates from humans and mice as representatives of the primate and rodent clades, respectively. We observe no difference in average *π* values between proteins evolving under positive selection versus purifying selection and/or neutral evolution in primates or rodents (Supplementary Figure 2A). Similarly, there is no significant correlation between dN/dS and *π* across kinetochore proteins (Supplementary Figure 2B). However, we cannot exclude the possibility that dN/dS estimates show alternative trends with diversity levels estimated from other rodent or primate species.

### Pervasive evolutionary rate covariation among kinetochore proteins

Physically interacting or co-functional proteins should evolve at similar rates across a phylogenetic tree (Clark, et al. 2012). We performed an analysis of evolutionary rate covariation to identify proteins that have experienced coordinated bouts of accelerated or decelerated evolution across our 17 profiled species, suggestive of their functional inter-dependence at a molecular level. Most kinetochore proteins exhibit higher evolutionary rate correlations than expected by chance (15,875 of 16,587 protein pairs; 95.7%; Supplementary Table 4), indicating widespread coordinated evolution of kinetochore proteins. Overall, proteins within the CCAN show significantly higher correlations and reduced Normalized Tree Distance compared to proteins that function in other kinetochore compartments (p < 0.05; Figure 4). Thus, on average, CCAN protein evolution is strongly constrained by the need to maintain molecular interactions with other proteins.

**Figure 4:**
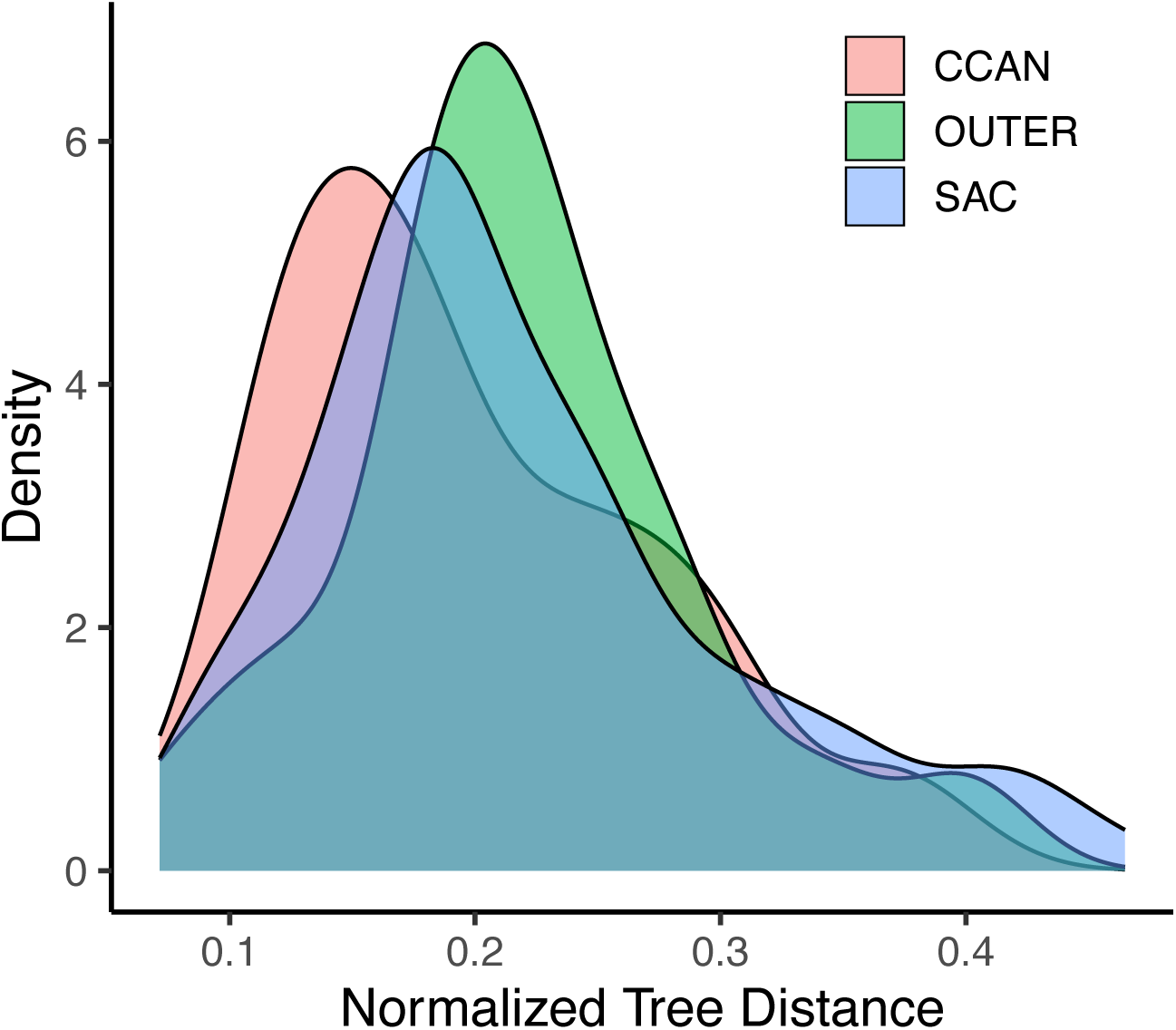
Density distributions of the Normalized Tree Distance across different classes of kinetochore proteins.

## DISCUSSION

Evolutionary theory predicts that the rapid evolution of centromere DNA satellite sequences should impose strong selective pressures on DNA-associated kinetochore proteins to maintain kinetochore function, preserve genome stability, and ensure accurate chromosome segregation. In support of this theory, prior investigations have documented signals of adaptative co-evolution between single kinetochore proteins and centromere satellite DNA (Henikoff, et al. 2001). However, the kinetochore is a large multiprotein complex comprised of a dynamic cast of structural, signaling, chromatin remodeling, and mechanical proteins. A single gene approach overlooks this complexity and is agnostic to the correlated evolutionary pressures experienced by physically associating proteins, including those than may not directly bind centromere DNA. Here, we have taken a holistic approach to comprehensively survey all protein components of the kinetochore. This strategy provides a cohesive, integrated portrait of kinetochore evolution and its relationship to centromere satellite sequence and architectural divergence.

Our analysis exposes three major trends. First, CCAN and outer kinetochore proteins evolve at similar rates. This result confirms reports from a prior study that examined the evolution of kinetochore proteins in primates, extending these findings to a wider group of mammals (Schueler, et al. 2010). Second, we demonstrate that many kinetochore proteins are adaptively evolving in concert with specific centromere features, including the presence/absence of a CENP-B box and HOR structure. Thus, beyond the linear DNA sequence of the centromere, aspects of the centromere chromatin environment may be important drivers of kinetochore protein evolution. Third, we observe correlated evolution among the majority of kinetochore proteins, with CCAN proteins exhibiting especially strong signals of evolutionary rate covariation. These results imply that adaptive changes in one kinetochore protein can often exert compensatory pressures on other proteins, effecting a “rippling” of adaptive evolutionary changes across the kinetochore complex.

Of the 215 kinetochore proteins profiled by our analyses, 52 (24.2%) exhibit signals of adaptive evolution in at least one phylogenetic test across rodents and primates. Thus, an appreciable proportion of centromere-associated kinetochore proteins are adaptively evolving as either a direct or indirect consequence of rapid centromere sequence evolution. Below, we profile several proteins harboring signals of adaptive evolution across Euarchontoglires and speculate on plausible selective pressures based on their known biological functions.

### Case studies in adaptive evolution across primates and mammals

#### Adaptive evolution in the CENP-A domain interacting with flexible DNA ends may protect CENP-A nucleosome integrity

Nucleosomes containing CENP-A, the centromere-specific histone H3 variant, form the scaffold for the assembly of the kinetochore complex (McKinley and Cheeseman 2016). In contrast to the remarkable conservation of histone H3, the amino acid sequence of CENP-A is highly variable between species (Henikoff, et al. 2001). Adaptive evolution in the N-terminal region of CENP-A has been previously observed in studies of eukaryotes (Henikoff, et al. 2001), *Drosophila* (Malik, et al. 2002), primates (Schueler, et al. 2010), and plants (Talbert, et al. 2004). However, studies profiling a wider range of mammals (Talbert, et al. 2004) and rodents (Kumon, et al. 2021) have not detected adaptive evolution at CENP-A. In our site model analysis of rodents and primates, we find evidence for adaptive evolution in the CENP-A domain encoding the flexible DNA ends that protrude from the nucleosome (Figure 5A). Whereas a canonical nucleosome contains ∼147bp of DNA, a human CENP-A nucleosome wraps only 121bp, leaving flexible DNA ends that are more vulnerable to nucleases (Kingston, et al. 2011; Tachiwana and Kurumizaka 2011) but nonetheless critical for the structural integrity of the CENP-A nucleosome (Roulland, et al. 2016). Rodent and primate species exhibit differences in centromere satellite repeat unit size, implying potential differences in the length and sequence of the flexible DNA ends associated with CENP-A-containing nucleosomes. Our finding raises the intriguing possibility that CENP-A evolves adaptively to ensure not only compatibility with centromere satellite sequence, but also centromere satellite length.

**Figure 5:**
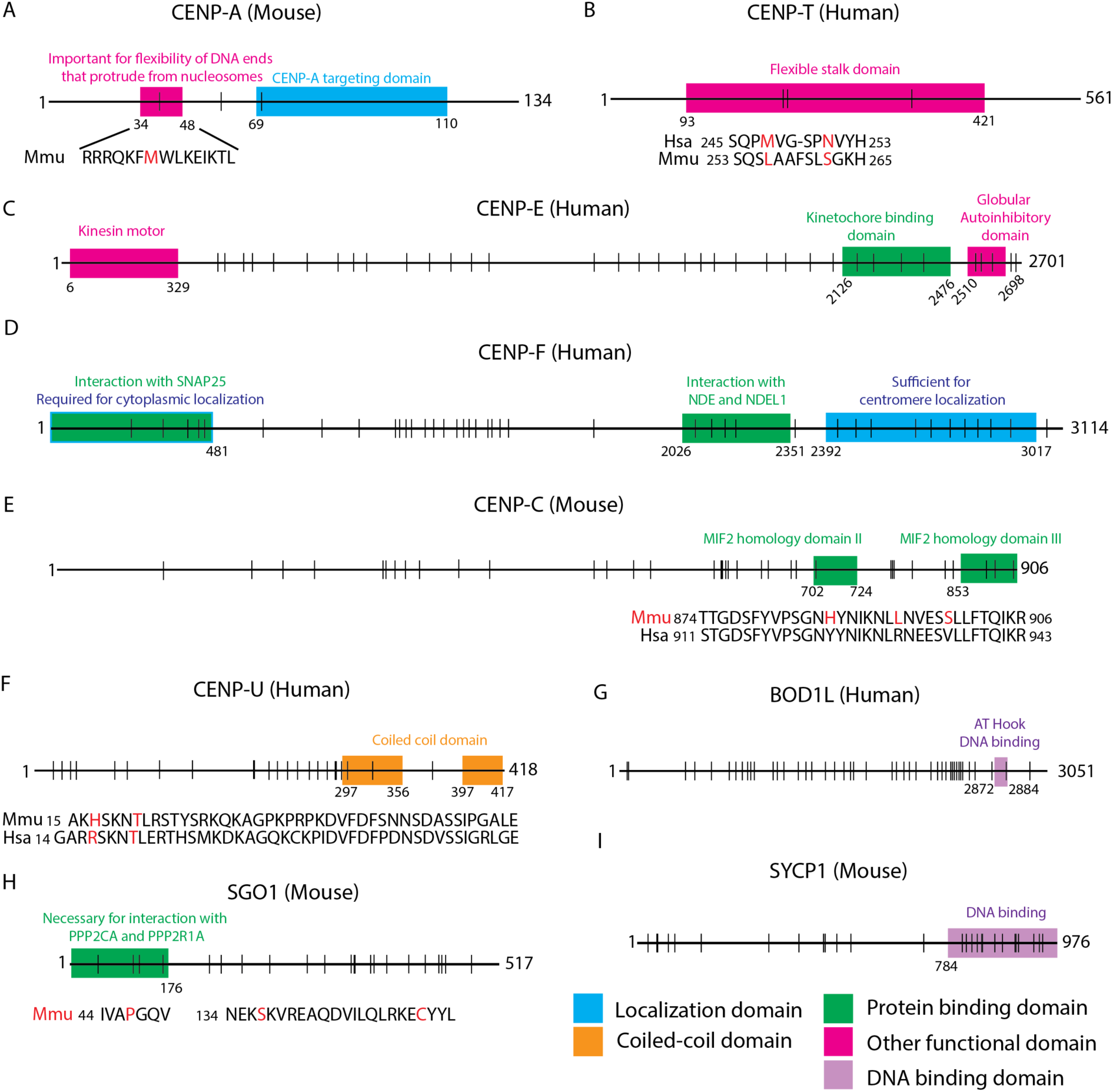
Candidate proteins under adaptive evolution. Panels A-I represent key kinetochore proteins identified in phylogenetic tests of adaptive evolution. Each protein is represented by a horizontal line, with colored boxes denoting known functional domains (legend on the bottom right) and vertical lines corresponding to positions of positively selected amino acids under the site and branch-site models. The species name in parenthesis represents the species from which the coordinates for the amino acids are depicted. Proteins with amino acid sites highlighted in red represent those under positive selection in the branch-site model. *Mus musculus* (Mmu) and *Homo sapiens* (Hsa) sequences are used as representatives for the rodent and primate groups respectively.

#### Adaptive evolution in CENP-T suggests modulation of CCAN assembly with centromere DNA variation

CENP-T forms part of the CENP-T/W/S/X complex that constitutively associates with centromere DNA (Nishino, et al. 2012). Specifically, the CENP-T/W heterodimer forms a histone fold domain that is essential for localization to centromere DNA (Takeuchi, et al. 2014). We identify three adaptively evolving sites in the flexible stalk domain of CENP-T (Figure 5B), adjacent to the globular domain conferring histone fold binding. Adaptive evolution in this domain could modulate histone fold binding in response to changes in centromere DNA sequence and chromatin structure between species.

#### Adaptive evolution of the outer kinetochore proteins CENP-E and CENP-F may stabilize kinetochore-microtubule binding dynamics

Accurate chromosome segregation depends on the efficient capture of kinetochores by microtubules and is facilitated by a suite of outer kinetochore proteins, including CENP-E and CENP-F (Cheeseman and Desai 2008; Nagpal and Fukagawa 2016). CENP-E and CENP-F exhibit inter-dependency in their localization; depletion of CENP-F results in a decrease in CENP-E, and vice versa (Johnson, et al. 2004; Yang, et al. 2005). Consistent with this inter-dependency, both proteins harbor signals of adaptive evolution in our site model analysis (Supplementary Table 1).

CENP-E is a microtubule plus-end directed kinesin motor protein that connects the kinetochore to microtubules from prometaphase through metaphase (Yen, et al. 1991). CENP-E facilitates chromosome congregation at the spindle equator (Putkey, et al. 2002; Kim, et al. 2008) and functions to actively maintain associations between kinetochores and dynamic microtubule ends (Gudimchuk, et al. 2013; Shrestha and Draviam 2013). Perturbation of CENP-E function results in the failure of kinetochores to form stable microtubule attachments, and ultimately leads to chromosome mis-segregation (Schaar, et al. 1997; Wood, et al. 1997; Yao, et al. 2000; Putkey, et al. 2002; Silk, et al. 2013). We identify adaptively evolving sites in the kinetochore protein binding domain of CENP-E (Figure 5D), raising the possibility that selection can modulate the strength of kinetochore-microtubule attachments.

CENP-F localizes to centromeres during prophase and to the kinetochore complex from prometaphase to anaphase. After anaphase, CENP-F remains in the spindle midzone and is rapidly degraded upon the completion of mitosis (Liao, et al. 1995). We identify sites under adaptive evolution within the centromere localization domain of CENP-F (Figure 5E), suggesting that changes in centromere satellite sequence can trigger co-evolutionary protein changes to maintain the CENP-F-centromere functional association. CENP-F localization at the kinetochore is dependent on BUB1 (Johnson, et al. 2004; Ciossani, et al. 2018), which is recruited to the centromere by the pericentromeric major satellite domain in mice (Akera, et al. 2019). Thus, evolution of pericentromeric repeat sequence and architecture could likewise impose selection pressures on CENP-F indirectly via BUB1. We also document amino acid sites under positive selection in the NDE1 and NDEL1 association domain of CENP-F. This domain is essential for correcting erroneous kinetochore-microtubule attachments (Auckland, et al. 2020). Whether mechanisms for correcting aberrant kinetochore-microtubule attachments vary across species and the potential extent of their dependency on centromere DNA sequence and structure remain open questions.

#### Adaptive evolution of CENP-C in rodents

CENP-C contains conserved domains involved in (1) binding of the MIS12 complex which associates with outer kinetochore KMN proteins (Przewloka, et al. 2011; Screpanti, et al. 2011) and (2) association with CENP-A resulting in the stabilization of CENP-A nucleosomes (Carroll, et al. 2010; Kato, et al. 2013; Falk, et al. 2015; Falk, et al. 2016). Together these conserved domains allow CENP-C to form a stable bridge between centromeric chromatin and microtubules, with depletion of CENP-C leading to chromosome mis-segregation (Falk, et al. 2015). Our findings replicate those of previous studies by uncovering signals of adaptive evolution of CENP-C in both rodents (Kumon, et al. 2021) and primates (Schueler, et al. 2010). Specifically, we document adaptively evolving sites in the MIF2 homology domain II and III domains at the C-terminus of CENP-C (Figure 5E). The MIF2 homology domain II is responsible for CENP-C association with CENP-A containing alpha satellite DNA, whereas the MIF2 homology domain III is important for self-dimerization in human cells (Trazzi, et al. 2009). The discovery of adaptively evolving sites in a region of CENP-C that appears to influence its association with centromeric chromatin (Carroll, et al. 2010) suggests that CENP-C, like CENP-A, may be locked in a co-evolutionary arms race with rapidly evolving centromere DNA.

#### Adaptive evolution of CENP-U in species with canonical CENP-B boxes

CENP-U is a CCAN protein that is part of the CENP-A nucleosome association complex (Foltz, et al. 2006). CENP-U is also a component of the CENP-O/P/Q/R/U complex; depletion of each of these complex components leads to mild mitotic defects *in vitro* (Hori, et al. 2008). However, during embryonic development, CENP-U function is essential, evidenced by the inviability of CENP-U null mice (Kagawa, et al. 2014). We identify adaptively evolving sites in CENP-U that are specific to lineages with centromere satellite repeats that contain canonical CENP-B boxes (Figure 5F). In *Mus musculus*, CENP-B protein modulates effector recruitment to destabilize microtubules, leading to preferential segregation of larger centromeres into the oocyte and provides a mechanistic basis for meiotic drive (Kumon, et al. 2021). We speculate that the coordinated evolution between CENP-U and CENP-B box status could provide a mechanism to counter altered CENP-B box abundance and supplement recruitment of kinetochore-microtubule regulator proteins. Indeed, CENP-U is known to recruit PLK1, a crucial effector protein that regulates kinetochore microtubule attachments (Chen, et al. 2021). However, it is also possible that CENP-U could influence CENP-B stabilization in the absence of CENP-B boxes. Further work is needed to test these hypotheses, and to ascertain the biological mechanism linking CENP-U adaptive evolution and CENP-B box abundance across species.

#### Adaptive evolution of BOD1L in species with higher order repeat centromere structure

BOD1L is involved in the replication fork protection pathway which functions to maintain genome stability in the face of replication stress (Liu, et al. 2016). Loss of BOD1L at stalled replication forks leads to excessive DNA resection and accumulation of DNA damage. Activity of the replication fork protection pathway is especially crucial for the maintenance of centromere sequence integrity. Centromere DNA can form non-B-form DNA structures which pose physical challenges for the replication machinery and lead to high rates of fork stalling (Black and Giunta 2018). We identified adaptive evolution in BOD1L at amino acid sites at and near the AT-hook DNA binding domain that presumably associates with centromere chromatin (Figure 5G). BOD1L dN/dS ratios are significantly different between species with and without centromere HOR structure (*ω* for species with HOR = 0.35, *ω* for species without HOR = 0.33, p = 0.004), suggesting that the dynamics of BOD1L association at stalled replication forks could vary across species in a manner dependent on centromere satellite architecture. Overall, our finding of selection signals uniquely associated with species harboring HORs suggests that complex repeat structures impose unique challenges for the faithful replication of DNA which may, in turn, impose selection on protein components of the replication fork protection pathway to increase replication fidelity.

#### Adaptive evolution of SGO1 in rodents

SGO1 is responsible for the maintenance of cohesin at centromeric DNA until deactivation of the spindle assembly checkpoint (SAC) and anaphase onset (Tang, et al. 2004; Watanabe and Kitajima 2005). This protective function is accomplished via recruitment of protein phosphate 2 (PP2) which protects cohesin subunits from phosphorylation and subsequent degradation (Clarke and Orr-Weaver 2006). We identify signals of adaptive evolution at sites in the PP2 interaction domain of SGO1 in the rodent clade (Figure 5H).

Prior work has established that the duration of female meiotic metaphase varies among *Mus* species (Akera, et al. 2019). Differences in the duration of meiotic metaphase can prolong or contract the timeframe over which stronger centromeres can detach from and reorient on the meiotic spindle to selfishly bias their segregation into the oocyte. At the same time, the length of metaphase defines the time window for operation of the SAC. Thus, the evolution of shorter metaphase may be a mechanism to counter centromere drive but may come at the expense of an increased rate of segregation errors due to reduced activity of the SAC. We speculate that adaptive evolution at the SGO1-PP2 protein interaction interface could be a mechanism for modulating metaphase duration via regulation of the timing of cohesion disassembly and anaphase onset, coordinating a balance between the competing pressures of minimizing the window of opportunity for selfish chromosome reorientation on the metaphase plate and ensuring the fidelity of genome transmission.

#### The DNA binding domain of the meiotic protein SYCP1 is adaptively evolving in rodents

SYCP1 is a component of the transverse filaments of the synaptonemal complex and is critical for ensuring pairing and crossing over between homologous chromosomes during meiotic prophase (de Vries, et al. 2005). We identified signals of adaptive evolution in SYCP1, reinforcing results from a previous study of meiotic protein evolution in mammals (Dapper and Payseur 2019). Although SYCP1 does not harbor a known sequence-specific DNA association pattern, we observe the highest rates of evolution in the DNA binding domain of SYCP1 (Figure 5I). This echoes observations for cohesin and condensin complex proteins, which are similarly sequence agnostic in their DNA binding, but are nonetheless adaptively evolving in mammals (King, et al. 2019). Centromeres are the site of initiation of synapsis during meiosis in budding yeast (Kemp, et al. 2004; Tsubouchi, et al. 2008) and *Drosophila* (Takeo, et al. 2011), but are the last regions to synapse during mouse meiosis (Qiao, et al. 2012). In mice, centromeres are also the last regions for synaptonemal complex disassembly, with SYCP1 functioning to stabilize bivalents at the metaphase plate and promote inter-centromeric connections that may facilitate proper homolog disjunction during meiosis (Qiao, et al. 2012). We speculate that SYCP1 may harbor sequence-specific binding for centromeric DNA or centromeric chromatin configuration, a possibility that would have likely escaped detection previously due to the gapped status of the centromere on most reference assemblies.

## Conclusions

We pursued a series of phylogenetic evolutionary analyses to investigate modes and mechanisms of kinetochore protein evolution. Our investigations show that the rapid evolution of centromere DNA has likely shaped rates of protein evolution across diverse kinetochore proteins, not just those that directly associate with centromeric chromatin. These findings support the hypothesis that rapidly evolving centromere DNA exerts a “ripple effect” across the kinetochore complex, wherein subtle changes at the protein-centromere DNA association interface can influence functional interactions between kinetochore proteins residing in distant physical domains of the large complex. As a consequence, organisms may be able to tune distinct kinetochore-mediated molecular pathways in response to changes in centromere DNA sequence, opening multiple avenues for preserving centromere function in the wake of rapid satellite sequence evolution and safeguarding against centromere drive.

In addition, we document multiple cases of adaptive protein evolution in species with specific centromere features. For example, we identify multiple kinetochore proteins with signals of positive selection restricted to species that carry centromere satellites with defined CENP-B boxes or limited to species with centromere repeats organized into HORs. Thus, extending beyond centromere DNA sequence evolution, changes in centromere architecture may also impose selective pressures on kinetochore proteins.

Taken together, our findings reinforce results from prior studies (Schueler, et al. 2010; van Hooff, et al. 2017), bolstering the sum of evidence for adaptive evolution at kinetochore proteins with diverse biological functions in microtubule attachment, DNA damage response, and chromosome cohesion. Importantly, our work also emphasizes the power of comprehensive investigations that interrogate the evolutionary history of all components of large protein complexes. When applied to other large protein complexes, this approach stands to uncover novel biology about the co-evolutionary dynamics of proteins connected through direct physical interaction and related through common biological networks.

## Materials and Methods

### Gene selection and codon sequence acquisition

We selected genes with GO term associations that contained the word “centromere” or “kinetochore”, yielding a set of 215 genes encompassing all known kinetochore proteins. We also included two histone H3 genes, which are known to evolve via purifying selection (Henikoff, et al. 2001). Codon sequences for each gene were downloaded from ENSEMBL (release 104). For each ortholog, we selected the transcript isoform that was the most highly expressed in most tissues.

### Species selection

We selected 17 members of the euarchontoglires mammalian clade encompassing rodents and primates with high-quality whole genome sequences: Human (*Homo sapiens;* Taxonomy ID 9606), Chimpanzee (*Pan troglodytes;* Taxonomy ID 9598), Gorilla (*Gorilla gorilla;* Taxonomy ID 9593), Bonobo (*Pan paniscus;* Taxonomy ID 9597), Orangutan (*Pongo abelii;* Taxonomy ID 9601), Macaque (*Macaca mulatta;* Taxonomy ID 9544), Marmoset (*Callithrix jacchus;* Taxonomy ID 9483), Mouse Lemur (*Microcebus murinus;* Taxonomy ID 30608), House mouse (*Mus musculus;* Taxonomy ID 10090), Steppe mouse (*Mus spicilegus;* Taxonomy ID 10103), Algerian mouse (*Mus spretus;* Taxonomy ID 10096), Ryukyu mouse (*Mus caroli;* Taxonomy ID 10089), Shrew mouse (*Mus pahari;* Taxonomy ID 10093), Norway rat (*Rattus norvegicus;* Taxonomy ID 10116), Naked mole rat (*Heterocephalus glaber;* Taxonomy ID 10181), Chinese Hamster (*Cricetulus griseus;* Taxonomy ID 10029), and Rabbit (*Oryctolagus cuniculus;* Taxonomy ID 9986). A cladogram reflecting current understanding of the evolutionary relationship among these 17 species was contructed using ape (5.6-2) in R (4.0.5).

### Codon alignment and alignment filtering

Codon sequences were aligned using probabilistic alignment kit (PRANK), a phylogeny aware, probabilistic multiple alignment program (Löytynoja 2014). Numerous multiple alignment programs are available, each with strengths and weaknesses (Pervez, et al. 2014). We selected PRANK on the basis that it employs an algorithm that treats insertions correctly and avoids over-estimation of the number of deletion events (Löytynoja 2014). A cladogram reflecting the known relationships among the 17 selected mammalian species was used as input to PRANK and branch lengths were estimated on this constrained topology for each gene based on the codon sequence alignment.

Analyses of adaptive evolution are susceptible to false positives due to alignment errors. Removing unreliable alignment regions is crucial for increasing the accuracy of evolutionary inference. However, this quality control step may significantly decrease the power to identify regions of adaptive evolution, as such regions are fast evolving and often difficult to align. Previous work has concluded that the benefits of removing unreliable regions outweigh the loss of power due to the removal of some true positive selected sites, and we adopt this conservative approach for our analysis (Privman, et al. 2012). We used default parameters in GUIDANCE2 to exclude sites with an alignment score below 0.93 (Penn, et al. 2010). We refer to these processed alignments as “filtered alignments”.

### Phylogenetic analysis of maximum likelihood (PAML)

Filtered alignments were input to the program Phylogenetic Analysis by Maximum Likelihood (PAML version 4.8) to measure the rate of evolution across the 17 species phylogeny (Yang 2007).

PAML measures the ratio of non-synonymous (dN) to synonymous (dS) amino acid substitutions in a maximum likelihood framework. PAML was run under 2 model frameworks for each gene: (i) the site model to assess which amino acid sites along a protein are under significant adaptive evolution and (ii) the branch-site model to identify the specific amino acid sites along one or more lineages that are evolving adaptively.

#### Site model

The site model was implemented under two different distribution frameworks –a maximum likelihood framework (M2 vs M1) and a maximum likelihood beta distribution framework (M8 vs M7). We note that the M2 model is named the M2a model in PAML v3.14 and earlier. The null models (M1 or M7) consider a fixed dN/dS values between 0 to 1 for codon sites along a gene. The alternative models (M2 and M8) allow dN/dS for a subset of codons to exceed 1. Codon frequencies were assumed to follow the F3x4 model of frequencies. We used the species tree for initial positive selection assessment, and subsequently used the gene tree output by PRANK (described above) as a confirmation. Nested models were compared using likelihood ratio tests under two degrees of freedom and a significance cutoff of 5.99 (corresponding to P < 0.05). For genes that had a significant likelihood ratio test under both models, Bayes Empirical Bayes (BEB) was used to calculate the posterior probabilities for site classes and identify sites under positive selection (i.e., codons within the gene that exhibited dN/dS greater than 1). We focused on sites under positive selection in functionally annotated domains.

#### Branch Site model

The branch-site models implemented in PAML can be used to detect evidence for positive selection affecting specific sites along prespecified foreground lineages. We implemented branch-site models per recent best practice recommendations (Álvarez-Carretero, et al. 2023), considering 4 different foreground branch designations (see below) applied to each kinetochore protein. The following null model parameters were specified: model = 2, NSsites = 2, fix_omega = 0. We invoked the following parameters for the alternative model, wherein a subset of sites is evolving either via relaxed selective constraint or positive selection across foreground branches: model = 2, NSsites = 2, fix_omega = 1, omega = 1. Models were compared using a likelihood ratio test, with 2 degrees of freedom and a significance cutoff of 5.99 (p < 0.05). As with the site model, BEB was used to calculate the posterior probabilities for site classes under the branch-site model and to identify sites under positive selection along lineages.

Species higher order repeat (HOR) status was assigned based on the designations of (Melters, et al. 2013). Species with HOR centromere structure include: *Rattus norvegicus*, *Homo sapien*s, *Pan paniscus*, *Pan troglodytes*, *Pongo abelii*, and *Microcebus murinus*.

The presence/absence of the CENP-B box within centromere satellite sequences of each species was determined from published studies. The canonical CENP-B box sequence is 17 bp (YTTCGTTGGAARCGGGA), but only 9 nucleotides are essential for CENP-B binding (N<u>TTCG</u>NNNN<u>A</u>NN<u>CGGG</u>N). Species with a high abundance of functional CENP-B motifs include: *Callithix jacchus*/Common Marmoset (Suntronpong, et al. 2016), *Homo sapien*/Human, *Pan paniscus/*Bonobo, *Pan troglodytes*/Chimpanzee, *Gorilla gorilla*/Gorilla, *Pongo abelii*/Sumatran Orangutan (Haaf, et al. 1995), *Mus musculus*/House mouse, and *Mus spretus*/Algerian mouse (Kipling, et al. 1995). The canonical CENP-B box for other species was assumed absent. Most centromeres in *Mus pahari* are comprised of repeats lacking the canonical CENP-B box, although the centromere of one chromosome carries an exceptionally high density of CENP-B containing repeats (Gambogi, et al. 2023). We group *Mus pahari* with species lacking CENP-B boxes.

### Nucleotide diversity

Nucleotide diversity (*π*) was computed from publicly available mouse (Harr, et al. 2016; Phifer-Rixey, et al. 2018) and human (Auton, et al. 2015) Variant Call Format (vcf) files (ftp.1000genomes.ebi.ac.uk, phase1_release_v3.20101123). Gene intervals for each of the 215 kinetochore proteins included in our analysis were extracted from University of California Santa Cruz Genome Browser and Mouse Genome Informatics database for human and mouse, respectively. Variants within each gene interval were extracted from the appropriate vcf file using bedtools (version 2.30.2). Nucleotide diversity (*π*) was computed by invoking the site-pi flag within vcftools (version 0.1.13) and converted to gene-level *π* values by averaging the *π* value across all sites in the focal gene. Nucleotide diversity for each group of kinetochore proteins was then Z-transformed to enable straight-forward comparisons across sets of functionally related genes. Mouse populations were analyzed by subspecies, with a total of 70 *M. musculus domesticus*, 22 *M. musculus musculus*, and 30 *M. musculus castaneus*.

### Evolutionary rate correlation

We assessed the evolutionary rate correlation between every pair of genes within the CCAN, Outer Kinetochore proteins (OUTER) and Spindle assembly checkpoint proteins (SAC). We used estimated trees from the Branch Site model for each protein, for which branch lengths were represented in units of number of substitutions/codon site. As necessary, each pair of trees was pruned to ensure perfect species agreement. Branch lengths were then rescaled so that the total length of each tree summed to 1. We then calculated the Pearson correlation coefficient between the scaled branch lengths of each pair of trees. To assess significance of the observed correlation, we generated a permutation-based P-value by randomizing the scaled branch lengths from one of the two trees and re-estimated the correlation on the permuted data. An empirical P-value was defined by the quantile position of the observed Pearson correlation coefficient along the distribution of 1000 permutation-derived values.

In parallel, we computed the normalized tree distance for each pair of gene trees, as previously described (Zheng, et al. 2015). Briefly, the normalized tree distance was calculated by summing the numerical differences between each pair of scaled branch lengths and dividing the sum by 2. Two identical trees will have a normalized tree distance of 0, whereas highly discordant trees will have a normalized tree distance close to one.

### Data availability

Raw data were derived from sources in the public domain: Ensembl (https://www.ensembl.org), UCSC Genome Browser (https://genome.ucsc.edu/), and Mouse Genome Informatics (https://www.informatics.jax.org/). Processed data are provided in supplementary tables 1-4.

## Acknowledgements and funding information

UA and BLD were involved in conceptualization, methodology, data interpretation, and writing, reviewing, and editing the manuscript. UA performed the formal analysis, and data visualization.

We thank members of the Dumont Lab for comments on the manuscript.

This work was funded by a Maximizing Investigators’ Research Award from the National Institute of General Medical Sciences to BLD (R35 GM133415). UA was supported by a Ruth L. Kirschtein Predoctoral Individual Fellowship from the National Cancer Institute (F31CA268727). The content of this manuscript is the sole responsibility of the authors and does not necessarily represent the official views of the National Institutes of Health.

---

**Supplementary Figure 1:**
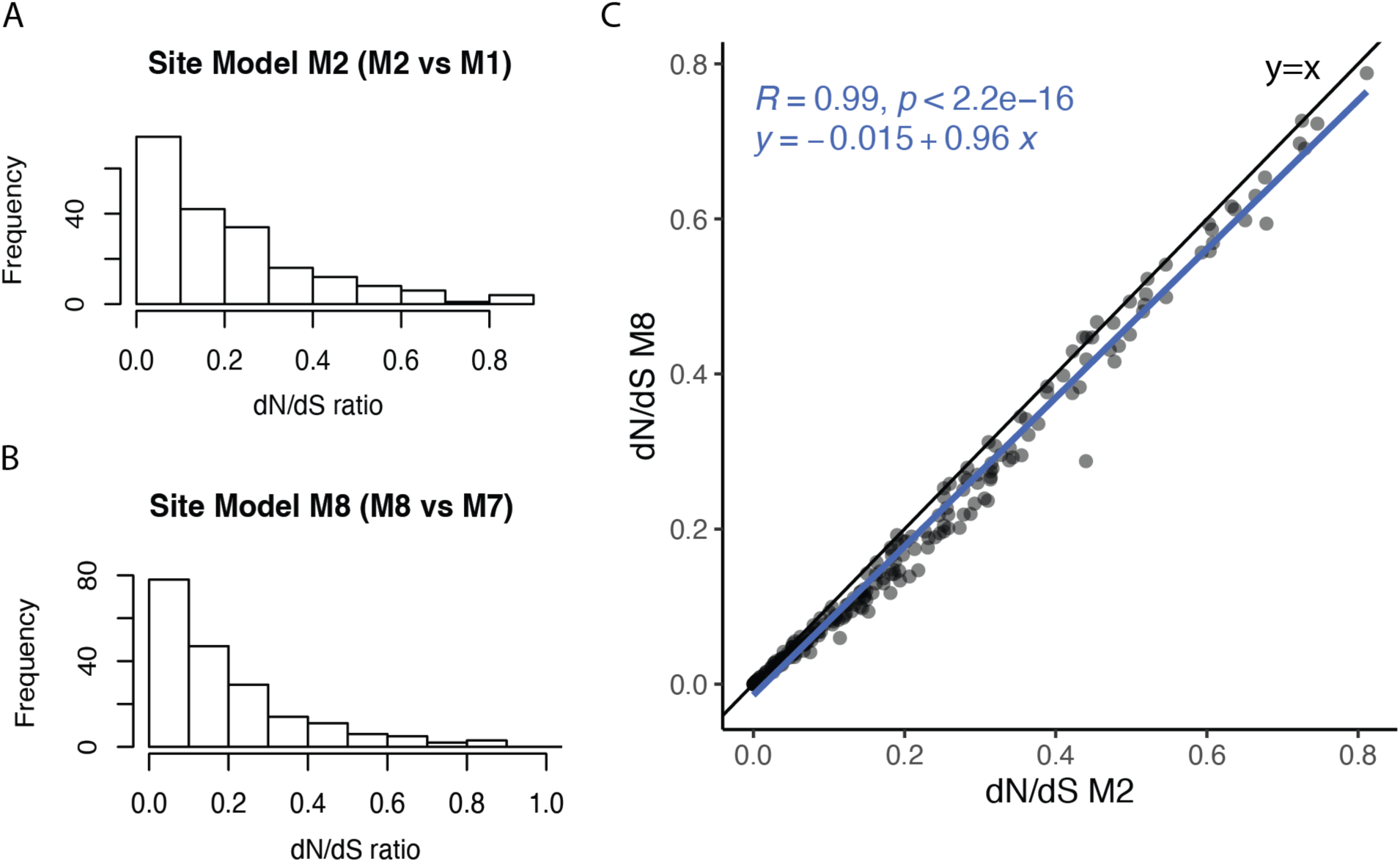
Majority of kinetochore proteins evolve under purifying selection. (A and B) Histograms representing dN/dS ratios (*ω*) of kinetochore proteins estimated under two implementations of the site model: (A) M2 and (B) M8. (C) dN/dS (*ω*) values under the M2 and M8 frameworks are strongly positively correlated.

**Supplementary Figure 2:**
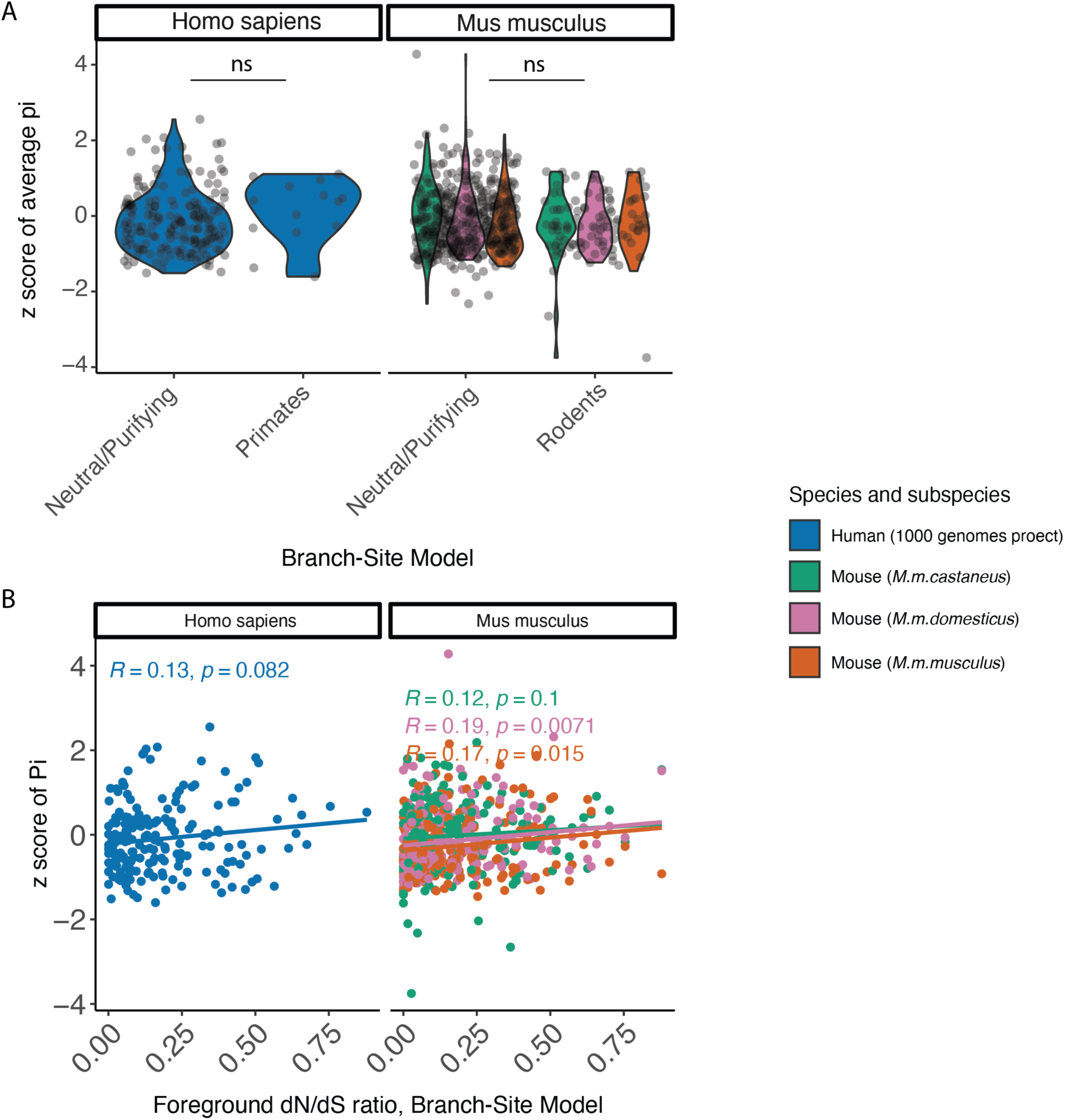
(A) Distribution of the normalized pairwise sequence divergence, *π*, across genes evolving via neutrality or purifying selection versus genes harboring signals of clade-specific adaptive protein evolution. The left panel features data from the primate branch-site model, whereas the right panel displays results from the rodent branch-site model. (B) Pearson correlation between the dN/dS estimated under the Branch-Site Model for each group (primates and rodents) and the normalized pairwise sequence divergence (*π*) in human and house mouse populations. Human *π* estimates were derived from 1000 Genomes Data. Mouse data are from Harr, et al. 2016.

